# Hierarchical Compression of *C. elegans* Locomotion Reveals Phenotypic Differences in the Organisation of Behaviour

**DOI:** 10.1101/029462

**Authors:** Alex Gomez-Marin, Greg J. Stephens, André E.X. Brown

## Abstract

Regularities in animal behaviour offer insight into the underlying organisational and functional principles of nervous systems and automated tracking provides the opportunity to extract features of behaviour directly from large-scale video data. Yet how to effectively analyse such behavioural data remains an open question. Here we explore whether a minimum description length principle can be exploited to identify meaningful behaviours and phenotypes. We apply a dictionary compression algorithm to behavioural sequences from the nematode worm *Caenorhabditis elegans* freely crawling on an agar plate both with and without food and during chemotaxis. We find that the motifs identified by the compression algorithm are rare but relevant for comparisons between worms in different environments, suggesting that hierarchical compression can be a useful step in behaviour analysis. We also use compressibility as a new quantitative phenotype and find that the behaviour of wild-isolated strains of *C. elegans* is more compressible than that of the laboratory strain N2 as well as the majority of mutant strains examined. Importantly, in distinction to more conventional phenotypes such as overall motor activity or aggregation behaviour, the increased compressibility of wild isolates is not explained by the loss of function of the gene *npr-1*, which suggests that erratic locomotion is a laboratory-derived trait with a novel genetic basis. Because hierarchical compression can be applied to any sequence, we anticipate that compressibility can offer insight into the organisation of behaviour in other animals including humans.

## Introduction

In introducing his four questions of ethology (1), Tinbergen emphasised that observation shapes how mechanistic and evolutionary questions are answered. That is, what we choose to measure determines the causal units that will form our explanations. The importance of understanding how animals structure their behaviour was recognised in part by the example set by genetics (2), in which many of the principles of inheritance were elucidated through careful observation and experimentation long before the physical nature of genes was known. For animal behaviour, the analogous goal is to generate or constrain hypotheses on the genetic and neural control of behaviour from the structure of behaviour itself.

Advances in automated imaging and computer vision make it possible to revisit the question of behavioural representation without relying on expert annotation. These methods have been used directly for quantitative phenotyping to measure behavioural differences in response to genetic and neural perturbation (3–11), as well as to study the dimensionality, dynamics, and structure of animal behaviour (12–16). However, even with the latest technology, automated analysis in complex natural environments remains challenging (17). Instead, we study the full structure and complexity of a behavioural repertoire in a simpler environment and focus on the spontaneous crawling of the nematode worm *C. elegans* confined to the two-dimensional surface of an agar plate. We have recently introduced a discrete representation of crawling postures and used it to identify short behavioural motifs that worms use to respond to sensory stimulation or that differ between worm strains isolated from different parts of the world (18). Here we explore whether data compression algorithms, which have been applied in domains where discrete data are common such as natural language processing and genomics, can reveal structure in worm locomotion.

Our approach is based on the minimum description length principle that the best model is the one that describes the data most concisely (19). We apply the minimum description length principle to behaviour by first constructing a dictionary of elementary behavioural states and then merging these states into longer sequences using a data-compression algorithm. The resulting new dictionary then serves as the ‘model’ of the behavioural data. Repeated steps of compression can find patterns and “patterns of patterns” in behaviour as proposed by Dawkins (2), thus generating a hierarchical representation of behavioural data. In addition, the degree to which these steps reduce the total length of the sequence and dictionary, the compressibility, offers a quantitative, objective measure of the behavioural complexity.

The connection between iterated dictionary compression and hierarchical organisation allows us to pursue two goals at once: to achieve maximum compression of the data and to mine its structure for biological meaning. In *C. elegans* we find that the dictionary sequences resulting from the compression algorithm represent rare but relevant behavioural motifs. We also measure the compressibility of behaviour and find that worm locomotion has intermediate compressibility poised between random and repetitive and that wild-isolates of *C. elegans* have locomotion that is more ordered than the laboratory reference strain N2 and mutants in an N2 background.

## Experiments

The data analysed in this paper comes from two previous studies (4,18). All N2, wild-isolate, and mutant tracking on food was done using single worms that were picked to the centre of a spot of *E. coli* OP50 on a 25 mm agar plate. Worms were allowed to habituate for 30 minutes before being tracked for 15 minutes. The worm side (whether it was on its left or right side) was manually annotated using a stereomicroscope before transferring plates to the tracking microscope. For off-food and chemotaxis experiments, worms were picked to the centre of 55 mm agar plates and recorded immediately. The attractant for chemotaxis experiments was 1 μl of benzaldehyde (diluted 1:100 in EtOH).

## Behavioural Analysis

### Posture discretization and time warping

The angles of the worm midlines were determined at 49 equally spaced points (16). The continuously varying skeleton angles were then discretized by matching the posture in each frame to its closest match in a set of 90 postural templates that were derived from wild-type N2 worms using *k*-means clustering (Fig. 1A). For details on the clustering and discretization, see (18). Because the motion of the worm between frames is often smaller than the difference between the 90 template postures, this procedure leads to the same template being fit in several consecutive frames. In order to recognize repeated behaviours performed at different speeds, we use a simple non-uniform time warping: repeats are removed from the posture sequences (for example, the sequence {1, 2, 3, 1, 1, 1, 4, 1} would be reduced to {1, 2, 3, 1, 4, 1}). Nevertheless, temporal information is not lost since we record the duration of each sequence template for subsequent analysis. In order to compare results across mutant strains and conditions, we used the same wild-type posture templates in all cases.

### Compression algorithm

Many popular dictionary compression algorithms are designed to work ‘online’ with little memory and to scan the data from left to right in a single pass looking for repeated patterns. We are more interested in finding repeating patterns than in single-pass memory-efficient compression and so we use an ‘offline’ algorithm that considers the entire sequence at each iteration. We follow Nevill-Manning and Witten’s offline ‘Compressive’ heuristic for inferring hierarchies of repetitions in sequences (20). This is a dictionary compression method in which repeated subsequences are added to a dictionary and replaced by a new symbol that indicates where the subsequence is contained in the dictionary. At each iteration, the subsequence that is replaced is the one that gives the maximal compression taking into account its length and frequency as well as the size of the dictionary. The savings, *S*, due to replacing a subsequence is given by *WN* ‐ (*W* + 1 + *N*), where *W* is the length of the subsequence and *N* is the number of times it occurs in the sequence that is being compressed. The first term is the reduction in the length of the sequence while the second term includes the increase in the size of the dictionary (*W* + 1) and the number of new symbols introduced in the compressed sequence (*N*). In the case of ties, where two subsequences are equally compressive, the subsequence that appears first in the sorted list of unique subsequences is replaced and added to the dictionary. This procedure is applied recursively until no more compressive repeats are found. Note that the compression algorithm is lossless. The original sequence can be exactly recovered using the compressed sequence and corresponding dictionary. The algorithm is also greedy, taking the locally most compressive sequence at each iteration and thus not guaranteed to find the globally most compressive dictionary. See Fig. S1 for an extended example explaining how the sequence in Fig. 1 is processed.

For faster computation, we calculate *S* only for subsequences up to length *W_max_*. We used *W_max_* = 10 for the results presented here. Increasing *W_max_* to 15 gave identical results in 97% of cases in a test of 200 worms and where results differed, the difference in compressibility was small (Fig. S2). The compressibility of a sequence of uncompressed length *l* is given by the sum of the savings *S* at each iteration divided by *l*.

## Results

### Hierarchical compression of posture sequences identifies behavioural structure

Dictionary based compression relies on an ability to identify repeated patterns in a symbolic sequence. Worm locomotion can be converted to such a symbolic sequence by representing the continuously varying worm body shape as a sequence of discrete postures (Fig. 1A). In this representation, the original skeleton (in black) is matched by its nearest neighbour posture in a set of 90 template postures (in blue) at each point in time. The templates themselves are determined using *k*-means clustering, with *k* = 90 postures chosen to capture most of the variance of worm shapes (~80%) without being overly complex (18). Approximately repeated behaviours can now be found simply by identifying repeated symbolic sequences, or *n*-grams. An *n*-gram is any subsequence of symbols of length *n*.

In a dictionary-based compression algorithm, a sequence is compressed by adding an *n*-gram to a dictionary and replacing each instance of that *n*-gram in the original sequence by a new 1-gram not previously present in the sequence. Maximal compression is achieved for *n*-grams that are both long and frequent. We call these maximally compressive patterns ‘c-grams’ to distinguish them from the larger set of *n*-grams they are drawn from. This is illustrated for a simple sequence with two symbols in Fig. 1B. In this example, we save 7 symbols in total since the original sequence was 19 symbols long, the compressed sequence is 3 and the dictionary contains a total of 9 symbols. The compressibility per symbol is therefore 7/19, or 37% of the original sequence length (see Fig. S1 for a more detailed explanation).

**Fig 1:**
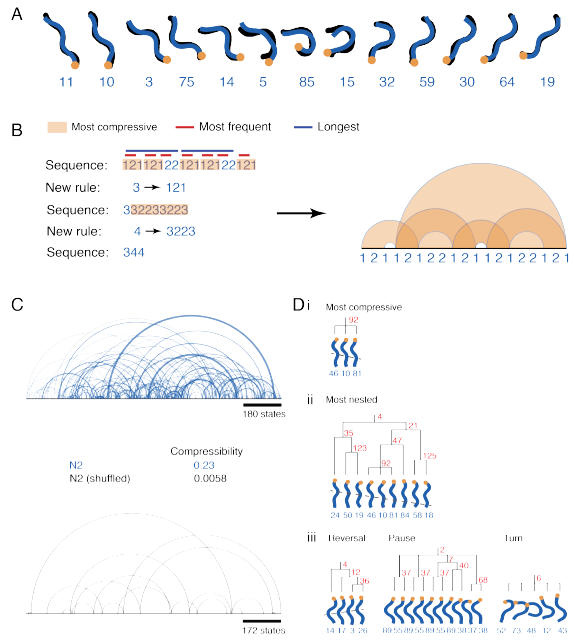
Dictionary-based compression extracts hierarchical structure in posture sequences. (A) Locomotion is represented as a sequence of discrete postural states. At each point in time, the original skeleton (black) is matched by its nearest neighbour posture in a set of 90 template postures. The orange dot indicates the head. The numbers beneath each shape are the labels of the template postures in each case. (B) Simple sequence to illustrate Compressive algorithm. For the indicated sequence, the subsequence that results in the greatest compression when it is replaced by a new state label is {1, 2, 1}. In the second iteration {3, 2, 2, 3} and {3, 3, 2, 2} are equally compressive. We simply take the sequence that occurs first in the sorted list of unique sequences. The arc diagram on the right connects adjacent repeats of dictionary sequences. (C) An arc diagram for a sequence of worm locomotion (blue) and the corresponding arc diagram for the same sequence following random shuffling (black). (D) Selected c-grams discovered from 150 minutes (~10^4^ postures) of worm behaviour. The most compressive sequence (i), the most nested c-gram (ii), and three other behaviours (iii) are plotted underneath dendrograms that show the hierarchical structure represented in the dictionary. The numbers in red indicate the number of times that the sequence under each branch occurred in the 150 minutes.

To visualize the replacement rules for a given sequence, we plot an arc diagram that connects each neighbouring c-gram that was used in constructing the dictionary (Fig. 1B, right). Frequently occurring c-grams are thus connected by short arcs, while rarely occurring behaviours are connected by longer arcs. The width of the arc corresponds to the length of the c-grams that are connected. When applied to wild-type worm locomotion (Fig. 1C), the arc diagram clearly shows that the majority of c-grams are frequent and short (small, thin arcs) but that there are some that are relatively rare and are separated by a long distance (longer arcs). The longest arcs connect c-grams that were only observed twice in the entire 1700 state sequence. This structure does not merely reflect chance repeats due to the finite number of symbols (the labels of the 90 template postures). When the sequence is randomly shuffled to maintain the posture frequencies but destroy temporal order, very few repeats are observed (Fig. 1C, bottom).

Notably absent in the arc diagrams of worm behaviour are long and highly nested repeats, which would be seen if worms perfectly repeated long sequences at different times. To provide further intuition for the level of repetition seen in spontaneous locomotion, we use the same algorithm to compress texts with increasing levels of structure and repetition: Moby Dick by Herman Melville (as used in a previous study on finding motifs in unannotated strings (21)), The Raven by Edgar Allan Poe, and Shake it Off by Taylor Swift (Fig. S3 and Fig. S4).

Some illustrative c-grams derived from a 30 minute (two 15 minute sequences concatenated) sample of wild type locomotion on food are shown in Fig. 1D. The most compressive sequence overall is shown at the top (i). This is the subsequence selected by the compression algorithm as providing maximum compression of the original sequence in the first iteration. Therefore, by construction, it is always “simple” in the sense that it has no nesting structure. In this case, it is a short bout of forward locomotion, consistent with expectations given that wild-type worms spend a significant portion of the time crawling forwards with a stereotyped gait.

The most nested c-gram found in the 30 minutes is shown in the middle (ii). Note that it contains the most compressive sequence from (i). Two suffixes are added (posture 84 appeared after the most compressive subsequence 47 times and this 4-gram was found with the 2-gram {58, 18} 21 times in a later iteration). Finally, a prefix completes the larger behavioural unit. This illustrates how units formed by basic templates are reused within larger units. In addition to various kinds of forward locomotion, we find c-grams corresponding to other behaviours. An example of a reversal, pause, and a turn are shown at the bottom of Fig. 1D (iii).

### Compressive sequences increase discriminative power across environmental conditions

Previous analysis has shown that the frequencies of 3-grams used by worms during locomotion can be used to characterize behavioural differences in different environments (on a lawn of bacterial food, on an agar plate without food, and during chemotaxis towards an attractant) (18). However, it is possible that sequences of other lengths are more informative for comparisons of behaviour in different conditions. The c-grams in the dictionary produced by hierarchical compression have variable lengths that are chosen adaptively based on the input data, in contrast to the fixed-length approach based on 3-grams. To determine whether they are relevant for behavioural comparisons, we reanalysed the data for worms in the different conditions.

We first compressed the postural sequences of each worm in each condition to produce a dictionary of c-grams for that individual. We then pooled all of the c-grams across all conditions, keeping only the unique c-grams, and compared the distributions of c-gram frequencies between conditions using rank sum tests, adjusted for multiple comparisons to control the false discovery rate at 5% using the Benjamini-Yekutieli procedure (22). Any sequence that was found to have a significantly different frequency between at least two conditions is a ‘hit’. The longest hits we detected were 10 postures long and represented bouts of forward locomotion. This was not because the maximum sequence length considered in a single iteration was 10 (see Behavioural Analysis above), since the same six sequences were still the longest hits when the maximum sequence length was increased to 15. One of the six 10-posture hits is shown in Fig. 2A and represents a persistent bout of forward locomotion which is most common during chemotaxis towards an attractant.

**Fig 2:**
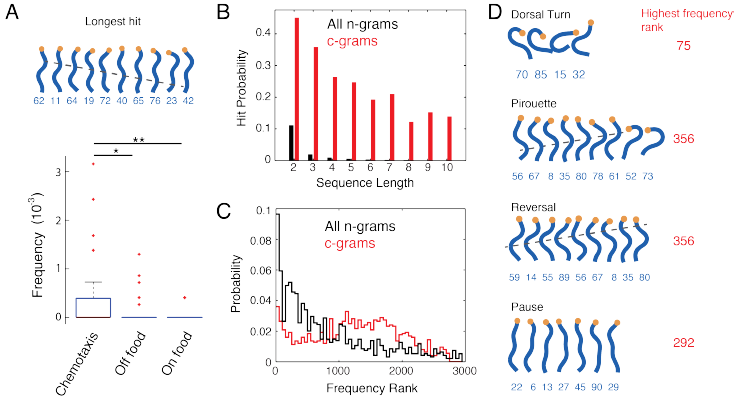
c-grams are rare but relevant subsequences. Hits are any sequences that are found to have a different frequency between N2 animals crawling on food, off food, or performing chemotaxis. (A) The longest hit is a bout of forward locomotion that is more common during chemotaxis. The box plot shows the frequency of this behaviour in the three conditions (red points are outliers, which are greater that the difference between the 25^th^ and 75^th^ percentiles outside of the box). (B) In each condition, the most compressive sequence is a hit in at least one comparison, indicating that compressive sequences are more likely to be modulated across conditions than n-grams as a whole. (C) The c-gram hits are more evenly spaced across the frequency distribution than those found using all *n*-grams. (D) Canonical worm behaviours are identified through compression and these would be missed by focusing only on the most frequently occurring *n*-grams. The behaviours are shown on the left with their highest frequency rank observed across all worms in the comparison group shown in red to the right.

The finding that c-grams with lengths up to 10 can be used to show behavioural modifications between conditions motivated us to revisit the previous analysis using *n*-grams with lengths of up to 10. For this relatively small data set consisting of 115 worms recorded for 15 minutes, this was tractable, but still required the consideration of 1.02 million unique *n*-grams. Of these, only 0.2% are used with a significantly different frequency in at least one of the conditions and this percentage is lowest for the longest sequences (Fig. 2B). In contrast, there were only 3014 unique c-grams in the entire pool, with 30% being significantly modulated in the environmental conditions. Furthermore, the fraction of significantly modulated behaviours remains high up to the maximum hit length. The *n*-gram hits are more likely to come from relatively frequent n-grams, whereas the c-gram hits are spread more evenly across the frequency spectrum (Fig. 2C). The precise frequencies and compressibilities change with the number of postures used in the representation, but the over all conclusions do not depend sensitively on the number of postures (Fig. S5).

It could be that the hierarchical compression algorithm is simply selecting frequent behaviours and that these are more likely to be informative for comparing worms in different conditions. To check this, we repeated the *n*-gram analysis, but took only the five most frequent *n*-grams of each length up to 10 from each worm and added it to the pool. This resulted in 3905 unique *n*-grams to use for comparing worms in the different conditions. This improved the efficiency of hit detection, in fact increasing it above that of the c-grams, especially for short *n*-grams (Fig. S6A). However, this improvement in efficiency comes at a cost: rare behaviours are no longer included in the analysis. This can be seen directly in the frequency rank distribution of the hits (Fig. S6B). Since this distribution includes the rank of each hit across all individuals, it is possible in principle that some of the *n*-grams that are among the 5 most frequent in one individual would be extremely rare in another individual, especially in a different condition. For the data considered here, that is not the case. The frequent *n*-gram distribution shows a much steeper drop-off than the c-gram distribution.

Examples of rare hits that would have been missed by focussing only on the most frequent n-grams are shown in Fig. 2D. These include potentially interesting behaviours such as a dorsal turn, a pirouette (reversal followed by turn), and a long reversal. Their highest rank across all individuals is shown in red above each behaviour.

### Worm behavioural sequences have intermediate compressibility

Hierarchical compression provides a new global feature for characterising worm behaviour: the compressibility of the sequences. It is clear that highly repetitive sequences will be more compressible than random sequences and we know from the plot in Fig. 1C that worm behavioural sequences are not random. We also know that compressibility must be greater than 0 and less than 1 by definition. To provide further intuition, we compared the compressibility of several ‘toy’ sequences (simulated controls) to real worm behavioural sequences as a function of sequence length (Fig. 3A).

**Fig 3:**
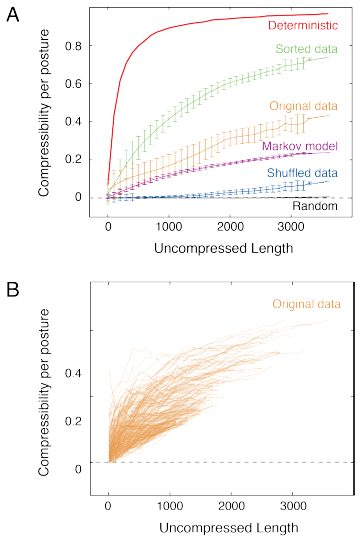
Worm locomotion sequences are poised between random and deterministic which leads to intermediate compressibility. (A) The compressibility per posture increases as a function of length for N2 locomotion sequences (orange). Uniform random sequences with 90 states (black) and a deterministic sequence consisting of 1 to 90 repeated (red) provide lower and upper bounds on compression. Shuffled (blue) and sorted (green) sequences provide related bounds constrained by having the same posture probability distributions as the observed locomotion sequences. A Markov chain simulated using the observed posture transition probabilities provides a more realistic model of locomotion sequences. (B) Compressibility as a function of length for individual worms shows the variability in compressibility. Many of the least compressible individuals have shorter uncompressed lengths, indicating that these worms moved less (had fewer posture transitions) during the 15 minutes they were recorded.

The first toy sequence we considered was a deterministic sequence which is simply the symbols 1 to 90 repeated in turn up to the desired length. This sequence is highly compressible, surpassing 0.8 compressibility for sequence lengths below 1000. Compressibility increases with length as more nearly optimal sequences are found, but can only reach 1 in the limit of infinite sequences. At the other extreme, we considered random sequences generated by sampling values from 1 to 90 from a uniform distribution. Uniform random sequences with 90 possible symbols are essentially incompressible for all observed lengths. At a length of 1000, the compressibility is 1 × 10^−4^ ± 3 × 10^−4^ (mean ± standard deviation). In contrast, behavioural sequences from wild type N2 worms crawling on food show intermediate compressibility, reaching 0.4 for the longest sequences considered. Control sequences that are more similar to real behaviour sequences were also generated by sorting and randomly shuffling behaviour sequences, yielding sequences that are more and less ordered than the original sequences but that have the same posture frequencies. Again, the natural sequences are poised between random and ordered. Finally, we also compared the natural sequences to sequences generated from a first order Markov model with transition probabilities determined from worm behaviour sequences. Although more similar than shuffled sequences, the Markov model sequences are still less compressible (i.e. less stereotyped) than the original worm sequences.

Plots of compressibility as a function of length for individual worms reveal inter-worm variability in compressibility (Fig. 3B). The least compressible worm sequences are also among the shortest, which result from worms that move less and therefore have fewer transitions. The fact that shorter sequences are more random suggests that the shape transitions that drive locomotion are more stereotyped than those that occur during dwelling. As expected, decreasing the number of postures in the representation increases compressibility (more repetition) and increasing the number of postures decreases compressibility (less repetition) (Fig. S7).

### Stereotypy varies across strains and does not simply reflect the degree of locomotion

Compressibility is a distinct feature for comparing the stereotypy of worm behaviour and so we analysed data from previously published mutant strains (4) and wild isolates (18).
Compressibility per symbol increases with length (Fig. 3) because there are more opportunities for compressive subsequences to be found in longer sequences. We therefore chose to compare worms using a fixed sequence length of 500. Sequences from worms that went through more than 500 distinct postures were divided into 500-posture chunks for analysis. Since dwelling worms seem to be less stereotyped than roaming worms (Fig. 3B) we also kept track of the time worms spent in each posture.

In Fig. 4A, we show a two-dimensional histogram of the distribution of compressibility against state duration for a set of 239 mutant strains that are not uncoordinated (‘Other mutants’). Each point in the histogram comes from one 500-posture chunk. The state duration value is simply the average time spent in each of the 500 postures. The overlaid lines are the full extent at half-maximum contours for the wild-isolate strains and for uncoordinated mutants. The distributions for the wild isolates and uncoordinated strains are plotted separately in Fig. 4B. Consistent with expectations from the N2 results, the wild isolate strains, which are known to move more persistently on food than N2 are highly compressible while worms that transition slowly between postures tend to be less compressible (there are few points in the upper right quadrant of the distribution). Nonetheless, differences in activity do not explain all of the variation in compressibility that is observed between strains.

**Fig 4:**
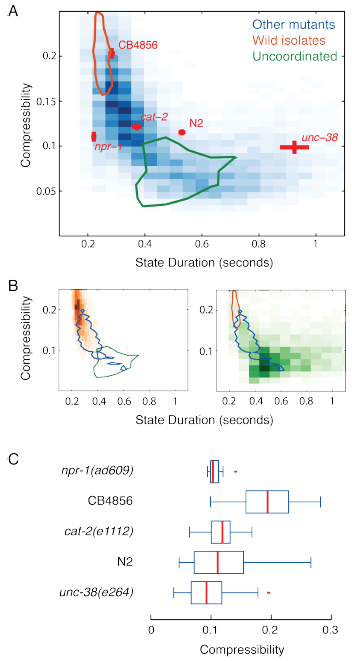
Wild isolate locomotion is more stereotyped than that of most mutant strains. (A) 2-dimensional histogram of the distribution of compressibility against postural state duration for a set of 239 mutant strains that are not uncoordinated (‘Other mutants’). The red bars show the mean ± standard error for a selection of strains. The contours show the extent at half-maximum of the distributions for 18 wild isolates (orange) and 63 uncoordinated mutants (green). The wild isolate and uncoordinated distributions are plotted separately in B. (C) Box plots showing the compressibility measured on 500 posture chunks for the strains highlighted in A. CB4856 is more compressible than either N2 (p = 4.7 × 10^−8^) or *npr-1(ad609)* (p = 3.3 × 10^−5^) using a rank sum test.

This variation is clear from the strains highlighted in Fig. 4A (red bars, mean ± standard error). The Hawaiian isolate CB4856 is known to be more active than N2 and it is also more compressible. However, two other hyperactive strains with loss of function mutations in *cat-2* and *npr-1* are significantly less compressible than CB4856 (Fig. 4C). This suggests that even though they move more persistently than N2, their locomotion is less stereotyped—more random—than that of CB4856.

These differences could be due to the use of N2-derived postures for all of the strains. This is a particular concern for uncoordinated strains that will adopt postures not seen in N2. We therefore re-derived postures for each of the strains individually and re-calculated their compressibility/duration histograms (Fig. S8). We also re-calculated the histograms using 250– and 1000–posture chunks (Fig. S8). The conclusions about relative compressibility are not altered in either case.

## Discussion

### Hierarchical structure in behaviour

The task of finding relevant behavioural motifs from a long string of postures is analogous to the task of finding genes in unannotated genomic data. However, unlike the situation in genomics, we do not yet have a ‘behavioural code’ that could guide the search. Instead, we take a more general heuristic approach to finding meaningful sequences inspired by the minimum description length principle. When we compress sequences of worm postures, we generate a hierarchical structure, but one that does not show a very high degree of nesting. Instead, the repeat structure of worm's spontaneous locomotion is characterised by short motifs that are used repeatedly but not normally in the identical context. In this sense, worms' spontaneous locomotion on food is more like a novel than a poem or song with a chorus (Fig S3). This is consistent with previous results using *n*-gram frequencies in worm locomotion. There is a small number of frequently used *n*-grams and a much larger set of rare *n*-grams (18). The structure we identify through compression suggests that the set of rare sequences is large enough to break up the repeated use of frequent patterns and to prevent the emergence of highly nested “patterns of patterns”.

A hierarchically organised action selection can lead to repetitive patterns in sequences (23), but the fact that a hierarchical representation can be constructed from a flat sequence does not necessarily imply that the underlying generating process is hierarchical. Instead, the nested structures we detect are best thought of as candidate behavioural units that may serve as hypothesised motifs for further study. Conversely, while there is more structure in worm behavioural sequences than in the corresponding shuffled data (Fig. 1C), we cannot rule out the presence of a deeper hierarchy in the underlying neural control. We would underestimate hierarchical structure if the output of a putative high-level command were implemented differently at the postural level because of environmental heterogeneity. That is, if different posture sequences were be used because of different local conditions despite the same overarching command.

The organisation of locomotion could be clarified by comparing patterns of behaviour with patterns of neural activity by imaging (24–28) and thermo-and optogenetic perturbation (15,24,29,30). Experimental manipulation of modular behavioural units was recently used to uncover a hierarchy of actions in grooming flies (23). A parallel model of action selection based on a suppression hierarchy was sufficient to reproduce the gross pattern of behaviours. In this case, the hierarchy of actions had a very simple structure in which activation of a higher behaviour suppressed the performance of the lower actions in the sequence. Dawkins referred to this kind of hierarchy as a ‘peck order’ (2) to distinguish it from more general control hierarchies that can have a complex branching structure. For complex hierarchies, even detecting the behavioural modules to probe may be more difficult. If modules can be identified and controlled, inferring the underlying control structure in the more complex case may be aided by using c-grams as candidate patterns to explore in more detail.

### Rare but relevant motifs

Compared to the total set of unique *n*-grams, the c-grams that are identified by hierarchical compression are a much smaller subset. In the case of worms in different environmental conditions, from the total set of unique *n*-grams only 0.3% were identified as c-grams. These proved to be a diverse set of behavioural motifs that were informative for comparisons between worms in different environments even though they were discovered on a per worm basis without reference to the environments the worms were in. Hierarchical compression can thus serve as a pre-processing step in behavioural comparisons that will make it possible to apply behavioural motif analysis to the large behavioural databases that are increasingly being created through high-throughput phenotyping pipelines (3,4,8,31).

### Compressibility as a quantitative phenotype

Compression provides a new measure for phenotyping that may give insight into mutant and wild-isolate differences. It was previously known that most wild isolates are faster on average than the laboratory strain N2, but we have found that this difference does not account for the differences we see in compressibility. For example, strains with a loss of function allele of the neuropeptide receptor gene *npr-1* show many of the phenotypes that are associated with wild isolates including increased speed, a shift in collective behaviour towards aggregation, as well as growth and pathogen avoidance (32,33). However, we find that *npr-1* mutant behaviour is less compressible than the wild isolate strains, including the well-studied Hawaiian strain CB4856. In other words, although they move persistently, their locomotion is more random than the wild isolates, as are the less persistent N2 worms.

The majority of mutant strains show patterns of locomotion that are less compressible (more random) than the wild isolates. Ranked in terms of compressibility, the 17 wild isolate strains have a median rank of 301 out of a total of 337 strains that were analysed and a maximum rank of 250. Compressibility is related to predictability, and being too predictable, especially in response to sensory stimulation can be deleterious in some circumstances; a fact that is strikingly demonstrated by tentacled snakes preying on fish (34). Unpredictability is also likely to be important for worms as recent work has demonstrated that ongoing network activity increases behavioural variability above the level predicted by sensory noise (35). Furthermore, a degree of randomness is an important element of *C. elegans* search strategies (36–38). Nonetheless, during directed locomotion, the most efficient gait is likely to be repetitive, and so we speculate that the high compressibility of wild isolates reflects a selective pressure for efficient locomotion and that the more random locomotion observed in N2 is due to a relaxation of this pressure in a laboratory environment. Laboratory domestication is known to have occurred in N2 based on the analysis of other phenotypes (39,40). Regardless of the ultimate cause, behavioural compressibility is a novel quantitative phenotype that is different between N2 and CB4856 that is not explained by loss of *npr-1* function. It therefore presents an opportunity to explore the genetics of this behavioural difference using recombinant inbred lines derived from these strains (41–45).

### Hierarchical compression beyond worms

Compressibility is a general measure that can be applied to the behaviour of any organism that can be tracked and discretised or converted to a series of labels by other means. The Nevill-Manning Compressive heuristic has already been applied to human motion capture data (46–48) and our approach could be readily applied to an ethogram derived either manually or automatically for any organism, including humans. This last possibility is worth considering because some human diseases affect locomotion (e.g. parkinson’s) and stereotypy (e.g. schizophrenia (49)) and compressibility might provide a simple scalar measure to quantify or even diagnose variation in a medically relevant phenotype.

## Acknowledgements

Some strains were provided by the CGC, which is funded by NIH Office of Research Infrastructure Programs (P40 0D010440).

## Data and code accessibility

The latest version of the code used for compression and to generate the figures in the paper can be found at https://github.com/aexbrown/Behavioural_Syntax. Worm crawling data are available at http://wormbehavior.mrc-lmb.cam.ac.uk/.

---

**Fig. S1:**
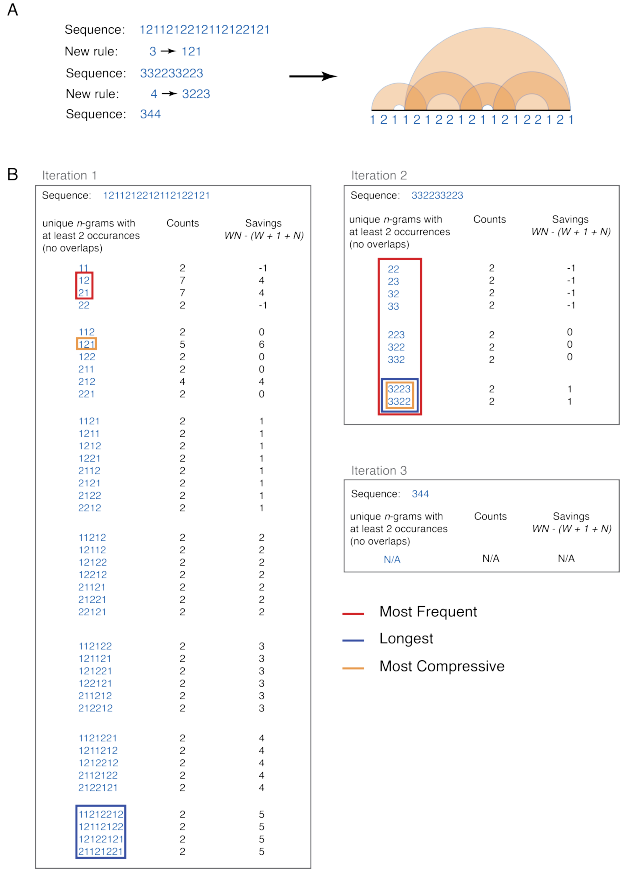
Sequences considered by a brute force version of the compression algorithm. (A) Summary of the algorithm and graphical representation of compressive sequences. (B) All nonoverlapping *n*-grams are counted. Only *n*-grams that occur at least twice in the sequence are included here (an *n*-gram that occurs only once can never compress the sequence because it takes as many characters in the dictionary as in the original sequence and requires the creation of a new symbol). Potential candidate sequences that would be replaced by different heuristics are highlighted, including the most frequent, longest, and most compressive. The counts and savings (total reduction in the number of characters after all occurrences in the original sequence are replaced with the new symbol) for each candidate sequence on shown. The savings can be negative which means that the size of the combined dictionary and compressed sequence will be larger than the uncompressed sequence. In the first iteration, there is a single sequence {1, 2, 1} which gives the maximum compression when replaced. However, ties are possible. In the second iteration, both {3, 2, 2, 3} and {3, 3, 2, 2} are tied for the most compressive sequence (in this case they are also the longest repeated *n*-grams). In the case of ties, we take the sequence that occurs first in the sorted list of *n*-grams, not necessarily the first to appear in the sequence, which is why {3, 2, 2, 3} is added to the dictionary in this case. The algorithm terminates when no further savings can be achieved, in this case that is because there are no more repeated *n*-grams, but it would also occur if there were only 3-grams with counts of 2 remaining since these would lead to zero savings when replaced.

**Fig. S2:**
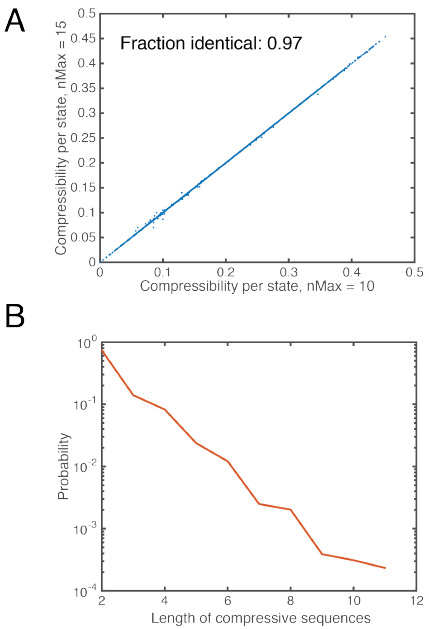
Increasing the maximum length of *n*-gram considered during compression has little effect on the observed compressibility. (A) For 200 worms each recorded for 15 minutes, the algorithm proceeded identically in 97% of cases. For those cases where a difference was observed, the difference was small. (B) The reason is that the probability of a sequence being the most compressive in a given iteration decreases exponentially with length. In the 200 worms, 11-posture compressive sequences were rare and 12-posture compressive sequences were not observed.

**Fig. S3:**
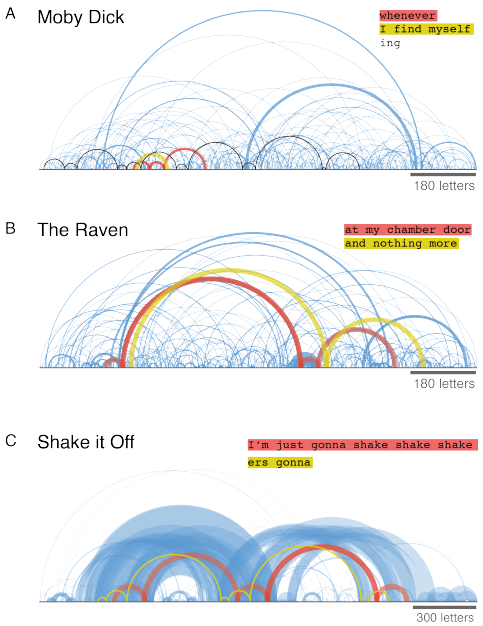
Arc diagrams show repeats in texts with increasing structure. In all cases, punctuation and spaces were removed and case was ignored. (A) The first 1200 characters of Moby Dick by Herman Melville. Prose includes some repetition, but usually only at the level of words and short phrases. Three uses of ‘whenever’ and two of ‘I find myself’ are highlighted. To find a more frequent repeat that occurs throughout, it is necessary to go to shorter structures such as the part of a word ‘ing’ which is shown as the black arcs. (B) The first 1200 characters of The Raven by Edgar Allan Poe. Wider arcs corresponding to longer repeats are visible. The relationship between ‘at my chamber door’ and ‘and nothing more’ are clear in the highlighted arcs. (C) Shake it Off by Taylor Swift shows strong repeat structure. The third use of the chorus is slightly different from the previous two and so the repeat is not perfect. ‘ers gonna’ is an interesting c-gram because it is used in two phrases: ‘Haters gonna hate’ and ‘Players gonna play’. The arc diagram shows that both uses have an invariant structure with respect to ‘I'm just gonna shake shake shake.”

**Fig. S4:**
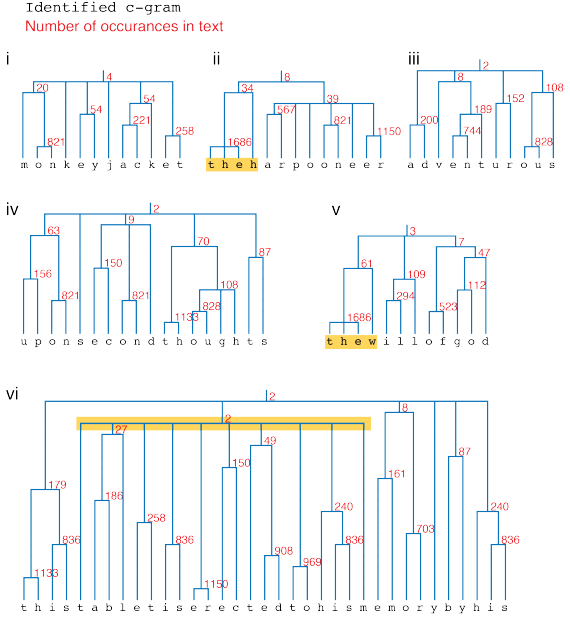
Repeated phrases found as c-grams in Herman Melville's Moby Dick are related to themes of the novel. The repeated phrases are related to the themes of the novel. The inferred hierarchy gives insight into the Compressive heuristic. Not surprisingly, 'the’ is found to be the most compressive sequence overall. However, because it is so frequent, it often gets associated with suffix letters and therefore breaks other words in the hierarchy. Two examples are highlighted in yellow in ii and iv. Because ‘will’ only occurs 34 times, 'thew’ is more compressive and is included in an earlier round of compression, effectively blocking the introduction of ‘will’ because of the greedy nature of the algorithm. iv is the longest repeat in the novel and illustrates a feature of language that is not commonly observed in worm locomotion. The highlighted branch of the dendrogram shows how 10 small branches merge in one step to form a phrase. While similar one-step merging of many short c-grams is observed in worm sequences (e.g. the pause in Fig. 1D), it is rare.

**Fig. S5:**
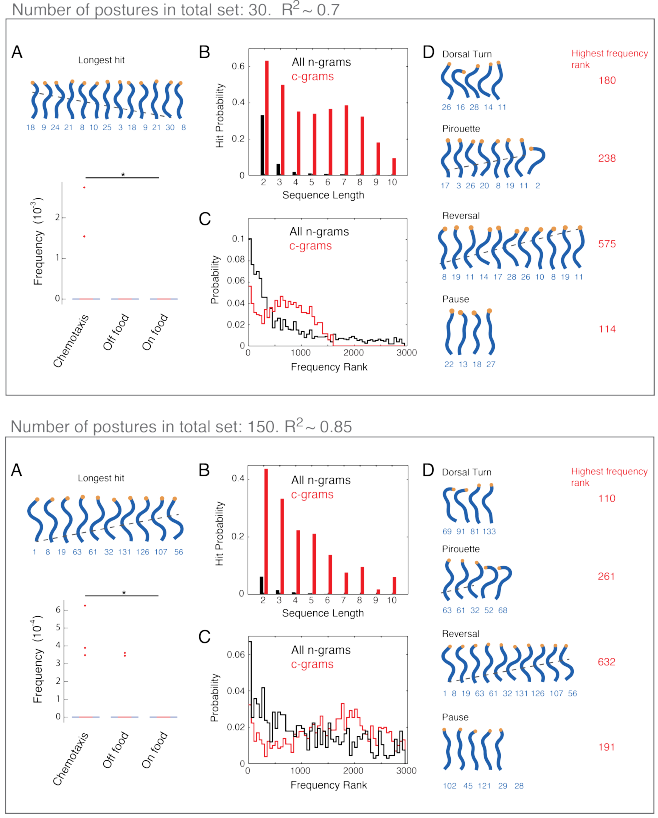
Changing the number of postures used to represent the worm behaviour changes the results quantitatively, but not qualitatively. The R^2^ value in gray for each posture number set indicates the average quality of the fit between the set of template postures and the original worm posture data from tracking. Reducing the number of postures leads to a less accurate fit but more compact representation and vice versa.

**Fig. S6:**
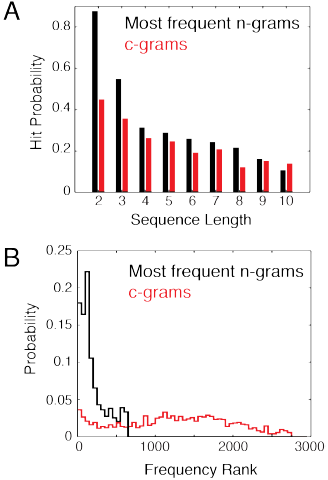
(A) The hit rate of *n*-grams is increased if only the 5 most frequent *n*-grams of each length from each worm are considered. However, this comes at the cost of ignoring rare sequences (B).

**Fig. S7:**
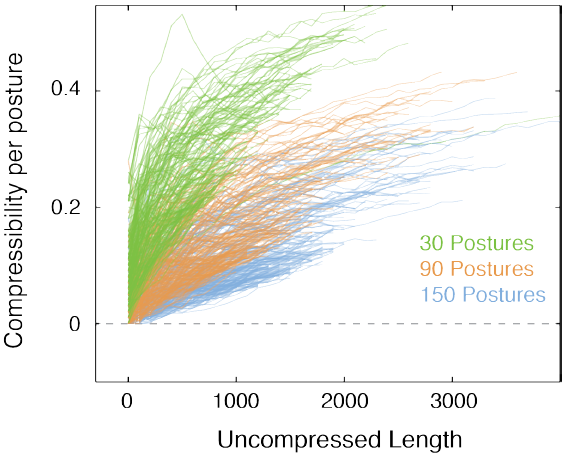
Increasing the total number of template postures used to represent behaviour decreases compressibility, in line with expectations.

**Fig. S8:**
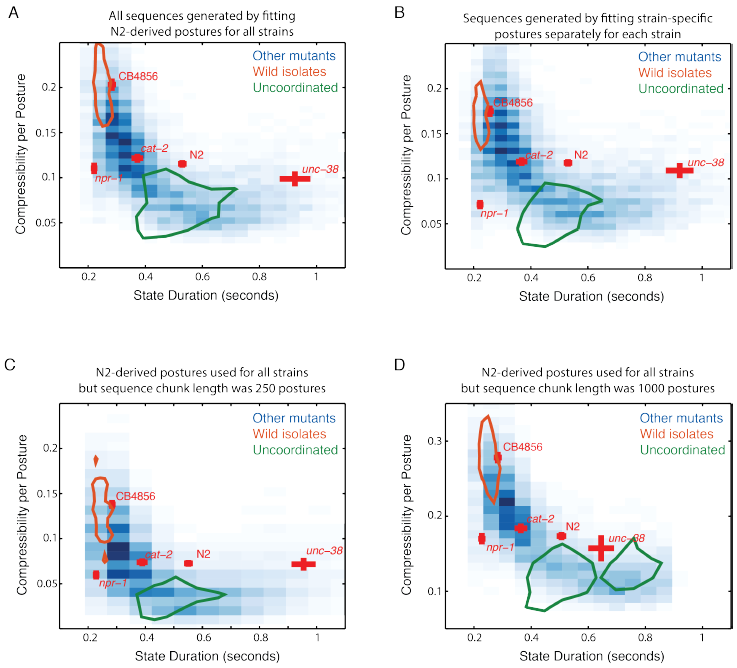
2-dimensional compressibility/state duration histograms. (A) The histogram using N2-derived postures and a chunk size of 500 postures, this is a reproduction of Fig. 4A in the main text. (B) The same histogram but using compressibilities calculated using template postures rederived for each strain. *npr-1* becomes less compressible in this case, but this only enhances the difference between it and the wild isolates. (C) This histogram was generated using N2-derived postures but shorter 250-posture chunks to calculate the compressibility. Compressibility is always lower, but the relationships between strains are similar. (D) Same as A and C, but using 1000-posture chunks for the calculation. Compressibility increases, but again the relative differences are similar. There is a truncation of the duration axis and a resulting shift in the *unc-38* duration because for slow-moving strains, some 15 minute videos include fewer than 1000 postures and were not included.

